# Spatially resolved integrative analysis of transcriptomic and metabolomic changes in tissue injury studies

**DOI:** 10.1101/2025.02.26.640381

**Authors:** Eleanor C Williams, Lovisa Franzén, Martina Olsson Lindvall, Gregory Hamm, Steven Oag, Muntasir Mamun Majumder, James Denholm, Azam Hamidinekoo, Javier Escudero Morlanes, Marco Vicari, Joakim Lundeberg, Laura Setyo, Aleksandr Zakirov, Jorrit J Hornberg, Marianna Stamou, Patrik L Ståhl, Anna Ollerstam, Jennifer Tan, Irina Mohorianu

## Abstract

Recent developments in spatially resolved -omics have enabled studies linking gene expression and metabolite levels to tissue morphology, offering new insights into biological pathways. By capturing multiple modalities on matched tissue sections, one can better probe how different biological entities interact in a spatially coordinated manner. However, such cross-modality integration presents experimental and computational challenges.

To align multimodal datasets into a shared coordinate system and facilitate enhanced integration and analysis, we propose *MAGPIE* (***M****ulti-modal **A**lignment of **G**enes and **P**eaks for **I**ntegrated **E**xploration*), a framework for co-registering spatially resolved transcriptomics, metabolomics, and tissue morphology from the same or consecutive sections.

We illustrate the generalisability and scalability of *MAGPIE* on spatial multi-omics data from multiple tissues, combining Visium with both MALDI and DESI mass spectrometry imaging. *MAGPIE* was also applied to newly generated multimodal datasets created using specialised experimental sampling strategy to characterise the metabolic and transcriptomic landscape in an *in vivo* model of drug-induced pulmonary fibrosis, to showcase the linking of small-molecule co-detection with endogenous responses in lung tissue.

*MAGPIE* highlights the refined resolution and increased interpretability of spatial multimodal analyses in studying tissue injury, particularly in pharmacological contexts, and offers a modular, accessible computational workflow for data integration.

## Main

Recent advances in spatially resolved -omics technologies have enabled untargeted studies of spatial distributions of various molecular species^1,2^. Spatially resolved transcriptomics (SRT) enables genome-wide spatial gene expression profiling^3–8^, allowing the untargeted exploration of disease-driving mechanisms at an unprecedented scale and resolution^9–13^. To further evaluate disease- and drug-induced alterations *in situ*, it is highly valuable to capture multimodal data^14^. The combination of SRT and mass spectrometry imaging (MSI)^15,16^ in the same tissue specimen has expanded the limits of spatial multimodal analyses^17–20^, facilitating the study of smaller, endogenous and/or exogenous molecules and metabolites, and opening new avenues for functional characterisation of local compound-induced tissue responses. While parallel unimodal analyses offer valuable insights, fully integrated spatial multi-omics workflows allow the direct investigation of cross-modality covariation within the same microenvironment and histological context. To unlock such analysis, it is essential to align all modalities into a shared coordinate system. However, generating multi-modal data for successful co-registration and co-analysis relies on both experimental and computational considerations.

On the experimental front, there are two established approaches: (1) developing protocols that allow multiple measurements on the same tissue section^18^ or (2) collecting each data modality on separate, consecutive tissue sections^17,19,20^. In both cases, sample preparation requirements ensure high analyte quality and compatibility with each modality. A typical approach for SRT would include optimal cutting temperature (OCT) tissue embedding; however, this obstructs MSI data collection^21,22^, prompting more appropriate methods to be considered for joint SRT and MSI analysis. Furthermore, refined methods may allow the addition of extra modalities for the assessment of tissue morphology, as captured by default in the Visium protocol and optionally obtainable after MSI data acquisition by histological staining and imaging of the tissue section.

Computationally, spatial analyte abundance levels and images obtained for each modality need to be co-registered to enable direct comparisons. Many current methods use ‘landmarks’ (corresponding locations across different images), identified either manually or automatically^23^. Once landmarks are identified from histological or spatial expression patterns, the geometric transformations required to align multiple images are estimated and applied. Given the inherent differences between consecutive sections and potential distortions introduced during sample preparation, straightforward linear mappings may fail to achieve accurate alignments and lead to compromised interpretations; more complex, non-linear, and elastic transformations may thus be required to effectively link corresponding spatial features across images. Several methods have recently been developed for this co-registration process, including SpatialData^24^ and Giotto Suite^25^, which offer purpose-built frameworks with data structures designed for spatial -omics, coupled with functions for alignment and aggregation of observations across modalities, including Visium and others. Nonetheless, these toolkits lack full interoperability with the existing landscape of spatial tools, such as R-based and multi-omics methods^26–29^ and further lack specific support for MSI data. SOmicsFusion^30^ supports MSI data, although it focuses on imaging data with no support for Visium datasets and is currently solely provided as standalone scripts, which may limit the ease of use for some users. STalign^31^ and Eggplant^32^ enable the alignment of multiple SRT datasets, however, they do not specifically support MSI data and have not been tested in the multi-omics setting. Across these existing methods, we noted a distinct lack of tools that perform standalone, accessible, and interoperable co-registration of spatial modalities, with particular support for combining sequencing-based and imaging-based modalities such as Visium and MSI. Such interoperability is essential to access existing and upcoming downstream analysis options that require matching spatial coordinates and observations^28,29,33–35^.

To overcome these challenges, we present a streamlined open-source computational workflow, ***MAGPIE*** (***M****ulti-modal **A**lignment of **G**enes and **P**eaks for **I**ntegrated **E**xploration*), for integrating spatial transcriptomics (Visium) and metabolomics (MSI) datasets, alongside histological images generated on same or consecutive sections. *MAGPIE* outputs standardised and readily usable files to be processed by downstream Python- or R-based spatial toolkits, making it versatile and scalable towards a wide variety of further computational spatial multi-omics analytical methods. The generalisability and scalability of *MAGPIE* was benchmarked on several datasets, covering different tissue types and MSI technologies. To illustrate the full versatility of the framework, we also generated and processed spatial multimodal datasets in two tissue injury studies using lung tissue. As the lung is an inherently heterogeneous tissue type, requiring specific care to preserve its integrity, we developed a new sample preparation strategy to augment compatibility with both SRT and MSI analyses.

In summary, *MAGPIE* showcases the capacity of efficient spatial multimodal data integration to reveal new insights into disease-driving mechanisms and to enhance the study of local drug-induced alterations within a tissue.

## Results

### Integration of spatially resolved transcriptomics and metabolomics with *MAGPIE*

To enable and benefit from full spatial multi-omics integration, each modality should be generated from the same tissue specimen. This may require appropriate adaptations of the experimental protocols for each modality and tissue type. Specifically, to enhance the quality and subsequent interpretability of spatial multi-omics datasets from Visium and MSI in lung tissue, a highly heterogeneous and fragile tissue type, we developed a new experimental sampling approach (**<u>Methods</u>**). Inflation of fresh rodent lung tissue using an agarose solution followed by direct freezing allowed preservation of tissue morphology while avoiding the need for OCT embedding, hence facilitating MSI analyses in tissue sections adjacent to those analysed with Visium to provide matched spatial multimodal data (**Fig. 1a**).

**Figure 1.**
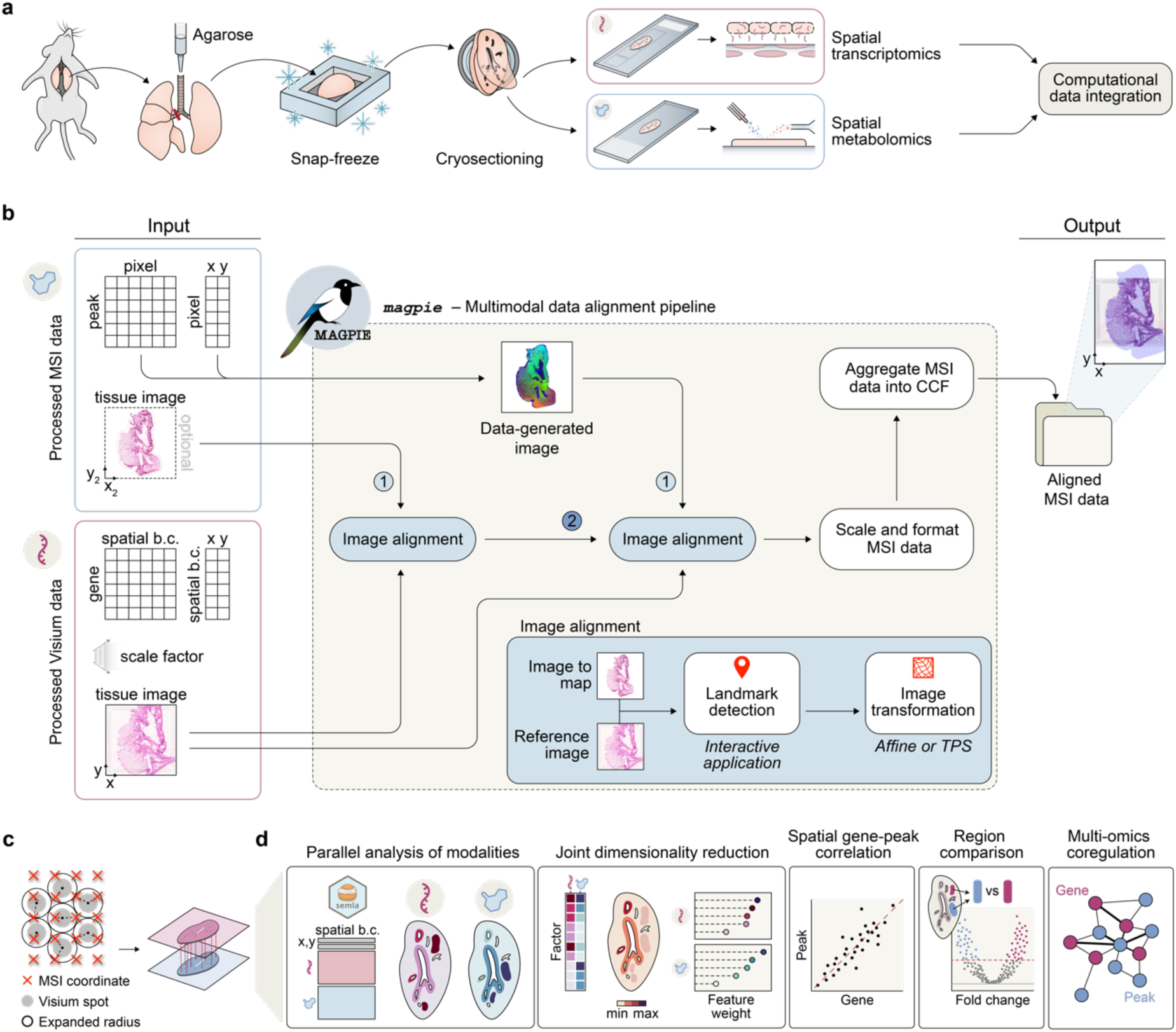
Spatial multimodal processing workflow and integration using the *MAGPIE* framework. **a**, Experimental workflow for spatial multi-omics profiling of rodent lung, consisting of agarose inflation of the lung tissue to facilitate the preservation of tissue integrity, followed by snap-freezing and cryo-sectioning to create consecutive sections, used for spatial transcriptomics and spatial metabolomics, respectively. **b**, *MAGPIE* computational framework for co-registering same or consecutive section spatial transcriptomics (Visium) and metabolomics (mass spectrometry imaging, MSI) data. The pipeline’s inputs and outputs are in standardised Space Ranger-tyle and tabular formats to ensure compatibility with other tools. Preprocessing the MSI data produces a data-generated image used for landmark selection and subsequent image co-registration. The output from the pipeline is the MSI data with updated coordinates aligned to the Visium data. **c**, To create a 1:1 mapping between Visium spots and MSI pixels, *MAGPIE* expands Visium spot radii, such that there are no gaps between spots, and then aggregates MSI pixels that fall within these spot boundaries. This results in matching observations between modalities and a processed object which can then be read by analysis toolkits, including *semla*^26^. **d**, Overview of downstream analysis options once the modalities are aligned into a matching coordinate system. Examples include joint clustering, dimensionality reduction, linking transcriptomic and metabolic changes to histologic information, and gene-peak correlation or multi-omics covariation network analysis. Abbrev. Spatial b.c., spatial barcode.

For the integrated computational analysis of spatial multi-omics assays, with a focus on transcriptomics and metabolomics, we developed the ***MAGPIE*** (***M****ulti-modal **A**lignment of **G**enes and **P**eaks for **I**ntegrated **E**xploration*) pipeline (**Fig. 1b**), implemented as a Snakemake workflow. As input, *MAGPIE* takes standard Space Ranger (10x Genomics) outputs from Visium acquisition and MSI data processed through standard software (e.g. SCiLS Lab or Cardinal^36^) into a tabular peak-by-pixel format. To co-register the MSI data with the image acquired from the Visium workflow, an MSI-representative image is generated in *MAGPIE* through dimensionality reduction of the MSI intensity data, allowing for the identification of tissue boundaries and morphological features. For this step, the user may guide the process to include *a priori* knowledge of spatially informative peaks or to colour each pixel based on 1-3 principal components (visualised as RGB channels). To support the alignment, the pipeline also accepts an intermediate tissue image for the MSI modality, such as a microscopy image of the tissue section, preferably stained similarly to the matching Visium section, e.g. with haematoxylin and eosin (H&E) or with immunofluorescence (IF). In this case, co-registration is first performed between the MSI data-generated image and the MSI tissue image, and thereafter between the MSI tissue image and the Visium tissue image. To manually select spatial landmarks for co-registration, *MAGPIE* includes an interactive (Python Shiny) application developed for this purpose. Alternatively, users can opt to incorporate externally identified landmarks, such as those generated using the deep learning-based unsupervised landmark detection method, ELD^23^, providing flexibility in the landmark selection process.

Once landmarks are identified, transformations of the queried MSI image to the reference Visium image can be performed using linear (e.g. affine) or non-linear (e.g. thin plate splines (TPS)) transforms. The transformation alignment projects the two modalities into a common coordinate framework (CCF), such that an (x,y) coordinate in one modality (MSI) can be directly mapped onto the other modality (Visium). Thereafter, the pipeline prepares the aligned second modality (MSI) data for outputting by converting all data into the standardised Visium Space Ranger-style output (i.e., MSI ‘count’ data in .h5 file format and coordinates, scale factors, and the transformed tissue image into a ‘spatial’ folder), which is readily accessible for downstream analysis using several spatial analysis ecosystems, such as *semla*^26^, Squidpy^37^, or others^24,27,38,39^. To perform a fully integrative analysis, pixels and spots are further combined into common ‘observations’, to ensure a matching at 1:1 resolution across the two modalities. To achieve this, *MAGPIE* maximally expands the radius of each Visium spot, while avoiding intersections with the neighbouring expanded spot radii, and thereafter aggregates (by sum or average) the values from all MSI pixels mapped within the expanded spot radius (**Fig. 1c**). The MSI pixel observations are thereby assigned to the same coordinate identifiers as used for the Visium spots. In addition, this aggregation approach has been added as a new function within the *semla* R package^26^ (v. ≥ 1.3.0) to allow the creation of a spatial multimodal object and further analysis within the Seurat or *semla* R ecosystems. Once fully integrated, a breadth of downstream spatial multimodal processing, analysis, and visualisation options are possible (**Fig. 1d**), facilitating novel discoveries on joint metabolomic and transcriptomic dynamics in healthy, diseased, or other perturbation settings.

### Data-driven guidance for enhanced *MAGPIE* co-registration

To ensure the successful co-registration of Visium and MSI datasets using *MAGPIE*, both technical and experimental factors need to be considered. In particular, we explored the effect of sample quality and parameter selection on co-registration performance, both of high importance when working with consecutive tissue sections where the section similarity is greatly affected by the sectioning process and tissue quality.

Several decision juncture points along the *MAGPIE* pipeline can impact the success of the resulting co-registration (**Fig. 2a**). Specifically, these user decision points are (i) whether a microscopy image for the second modality is generated or if co-registration is performed directly from the data-generated image, where successful landmark identification can be more difficult, (ii) the colouring used for the data-generated image, which may impact the ease and accuracy of landmark identification, (iii) how many landmarks are used and (iv) whether an affine or TPS transformation is used to map between modalities, of particular importance when the tissue sections used for each modality differ strongly. To assess these factors and guide the study design and parameter settings when running *MAGPIE*, we generated DESI MSI data and performed H&E staining of consecutive sections of mouse lung tissue (n=9) previously analysed with Visium^13^.

**Figure 2.**
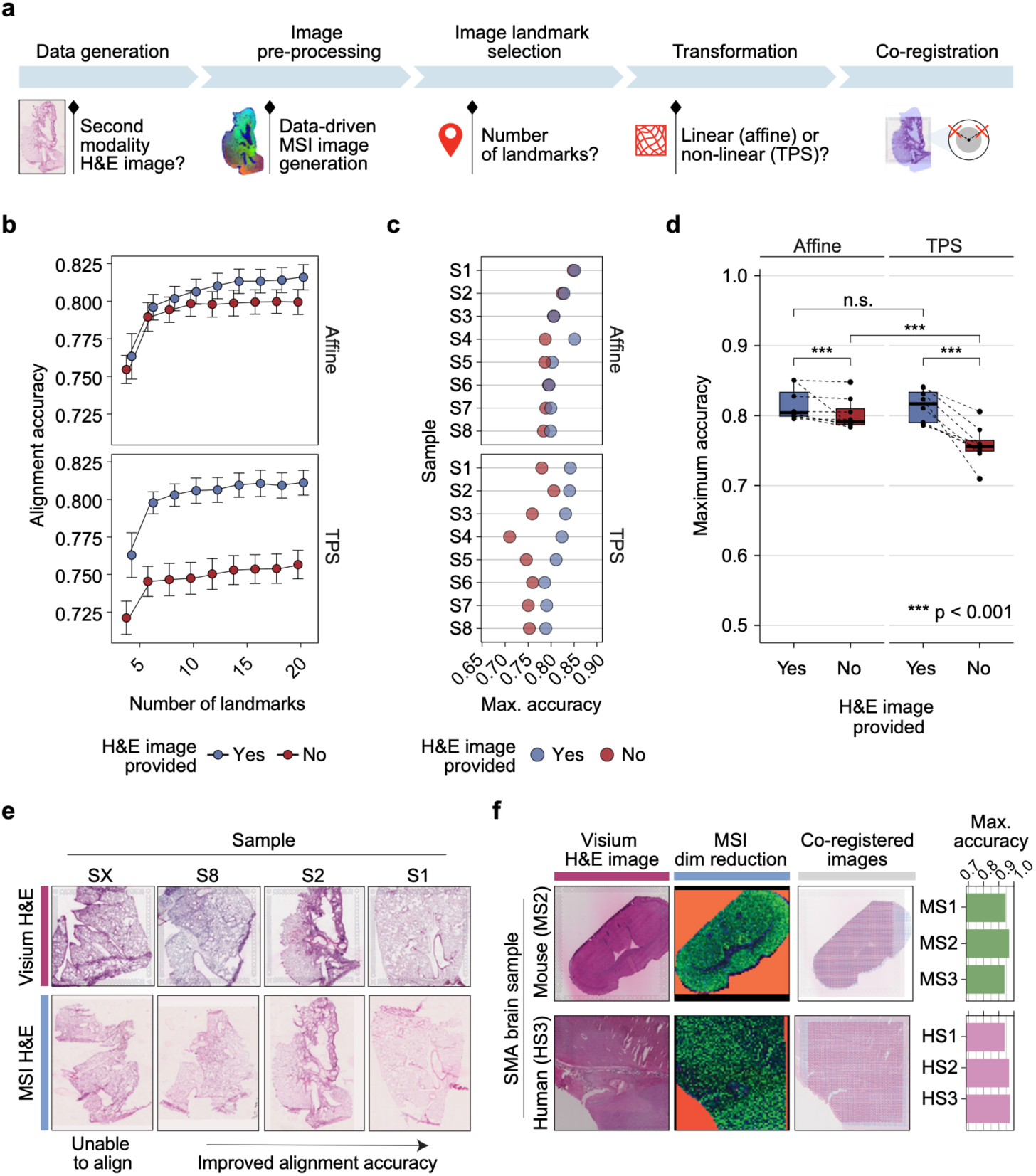
Demonstration of *MAGPIE* flexibility and robustness. **a,** Summary of the *MAGPIE* pipeline key decision points for user consideration. **b,** Variation in the alignment accuracy across a range of landmarks, employing a linear (affine) or non-linear (TPS) transform, with or without using an intermediate MSI H&E image to assist with co-registration, for 8 mouse lung samples. For each setting, the mean alignment accuracy per sample was selected across 5 iterations of randomly selected landmarks followed by averaging the scores across all samples, shown as points in the graph. Error bars correspond to the standard error. Points are coloured by the inclusion of an intermediate H&E image for the MSI modality. N samples = 8. **c,** Maximum accuracy for each sample using either affine or TPS transform, with or without the aid of an intermediate MSI H&E image. **d**, Comparison of highest accuracy performance per sample using affine or TPS transforms, with or without an intermediate H&E microscopy image. Each point represents the maximum achieved accuracy for each tested sample (n=8) with the boxplot showing the median, upper and lower quartiles with whiskers extending to 1.5× interquartile range. The maximum alignment accuracies per group were compared using paired two-sided Wilcoxon rank sum tests (Affine with H&E vs Affine without H&E: p = 1.12×10^-4^, TPS with H&E vs TPS without H&E: p = 1.77×10^-13^, Affine with H&E vs TPS with H&E: p = 0.198 (n.s., non-significant), Affine without H&E vs TPS without H&E: p = 1.69×10^-13^). **e**, Examples of H&E-stained tissue sections of mouse lung section pairs, ordered by maximum accuracy measurement between Visium and MSI sections. **f,** Generalisability of *MAGPIE* to other species and tissue types by application to same-section Visium and MALDI-MSI data (“SMA”) from mouse and human brain tissue. The maximum co-registration accuracies achieved per sample (mouse, n=3; human, n=3) are shown as bar charts.

Following the standard pre-processing of each modality for our generated paired Visium and MSI data, the data was processed with *MAGPIE* to benchmark the impact of altered input parameters. To assess the extent of co-registration success, we computed an alignment accuracy score based on the overlap between tissue and background labels in the Visium and MSI observations, specifically by comparing the number of spots labelled consistently in both modalities after transformation to the total number of spots (**<u>Methods</u>**). For the successfully co-registered samples (8 out of 11), up to 20 distinct landmarks per sample were manually selected using the interactive application within *MAGPIE*, allowing the assessment of alignment accuracy as a function of the number of landmarks (**Fig. 2b, Supplementary Fig. 1a**). Regardless of transformation type or the inclusion of an intermediate H&E image for the MSI modality, the accuracy score started to plateau at around 10 landmarks, with small additional improvements for some samples when including more landmarks (**Supplementary Fig. 1a**). To better understand the differences in co-registration between samples, we compared the maximum achieved alignment accuracy, across up to 20 landmarks, between samples with or without an intermediate H&E image and across image transformation types (affine vs TPS) (**Fig. 2c**). Compared to direct co-registration from MSI data to Visium, co-registration using an intermediate MSI H&E image attained higher alignment accuracy in terms of tissue overlap, with a larger difference observed when a TPS transformation was applied (**Fig. 2d**).

The co-registration accuracy differences across samples (**Fig. 2c**) underline extensive discrepancies between the sample section pairs, which may be explainable by the morphological distortions present within the H&E-stained tissue sections (**Fig. 2e**). The samples with the highest maximum accuracy were characterised by well-preserved tissue integrity, with only minor tears and folds **(Supplementary Fig. 1b)**. In cases where a section was too severely damaged, co-registration using manual landmarks was not possible (e.g. sample “SX”), while relatively high alignment accuracies could be achieved with *MAGPIE* where there were moderate tears and shape differences in the section (e.g. sample “S8”). Some of the more dissimilar samples demonstrated greater accuracy differences when intermediate MSI H&E images were utilised, highlighting their added value.

Based on these experiments and benchmarking results, we strongly recommended obtaining an intermediate MSI image (e.g. H&E staining after MSI) for more robust landmark identification and enhanced co-registration. In addition, for cases where landmark identification is challenging, opting for a linear transformation, such as affine, can help to avoid exaggerated distortion of the image transformation potentially caused by improperly placed landmarks. Conversely, for more dissimilar serial sections, a non-linear transformation may be necessary to achieve high-quality co-registration. Finally, an added benefit of selecting a high number of landmarks to aid the co-registration was observed, and we, therefore, recommend that users aim for higher numbers of manually selected landmarks.

### *MAGPIE* provides flexibility across tissue types and technologies

Next, we showcased the versatility of *MAGPIE* on samples from different species, tissue types, and MSI technologies. Co-registration was performed on previously published same-section Visium and MALDI MSI data from human (n=3) and mouse (n=3) brain tissue^18^ (Spatial Multimodal Analysis or “SMA” dataset) (**Fig. 2f, Supplementary Fig. 1c).** Testing up to 15 landmarks for each sample, all the included SMA samples achieved near-perfect co-registration accuracy (≥ 0.9), illustrating the successful alignment of datasets and the benefit of having same-section multimodal data (**Fig. 2f, Supplementary Fig. 1d**). Nonetheless, challenges in landmark identification and alignment accuracy assessment were encountered for the human brain SMA samples, which likely stemmed from their complex underlying structure and morphology and the lack of visible tissue boundaries, since the data was acquired from small regions of interest within a larger tissue block. Conversely, the mouse brain SMA samples had distinct morphological features and section edges that eased the manual landmark selection process. Hence, as a consideration, the placement and integrity of the tissue section can aid landmark selection if more distinguishable features are present, where holes and tears may even be beneficial. In addition, the choice of alignment metric can impact the presented co-registration accuracy scores, which in our case was strongly influenced by the tissue-to-background selection and differences between MSI and Visium capture area sizes. Focusing on image-level differences, rather than spot- or pixel-level, might present a complementary alignment accuracy score metric, albeit reliant on having an intermediate H&E image from the MSI modality, which was not available in our case.

Our results show that successful co-registration with *MAGPIE* is achievable for different tissue types and data acquisition technologies. Further, we highlight the importance of carefully selecting samples for multi-omics integration, particularly regarding the clarity of histological features, which are crucial for successful co-registration and especially important when working with larger or more complex tissue sections.

### Alignment of spatial multi-omics in pulmonary fibrosis reveals disease-associated signatures

To explore the insights into disease-driving mechanisms provided by *MAGPIE*-integrated spatial multi-modal data, we further analysed the dataset comprising Visium and DESI MSI on mouse lung samples (introduced in Fig. 2). These samples were collected from a mouse model of bleomycin (BLM)-induced pulmonary fibrosis, a widely used preclinical animal model for studying lung fibrosis and for the identification and assessment of new therapeutic drugs for human idiopathic pulmonary fibrosis (IPF)^40,41^. The administration of the cytotoxic agent BLM induces acute inflammation and subsequent fibrotic scarring of the lungs^42,43^. Recently, an in-depth Visium-based characterisation of tissue samples from IPF patients and the BLM mouse model was published, describing and comparing molecular signatures associated with the fibrotic niches^13^. To further build on this study and previous reports of metabolic alterations seen in IPF and BLM-induced pulmonary fibrosis^44–46^, our spatial multi-modal Visium and DESI MSI data from mouse lungs collected 21 days following BLM administration (n=6) were integrated and analysed within their histopathological context along with cell type mapping inference^13,47^ (**Fig. 3a**).

**Figure 3.**
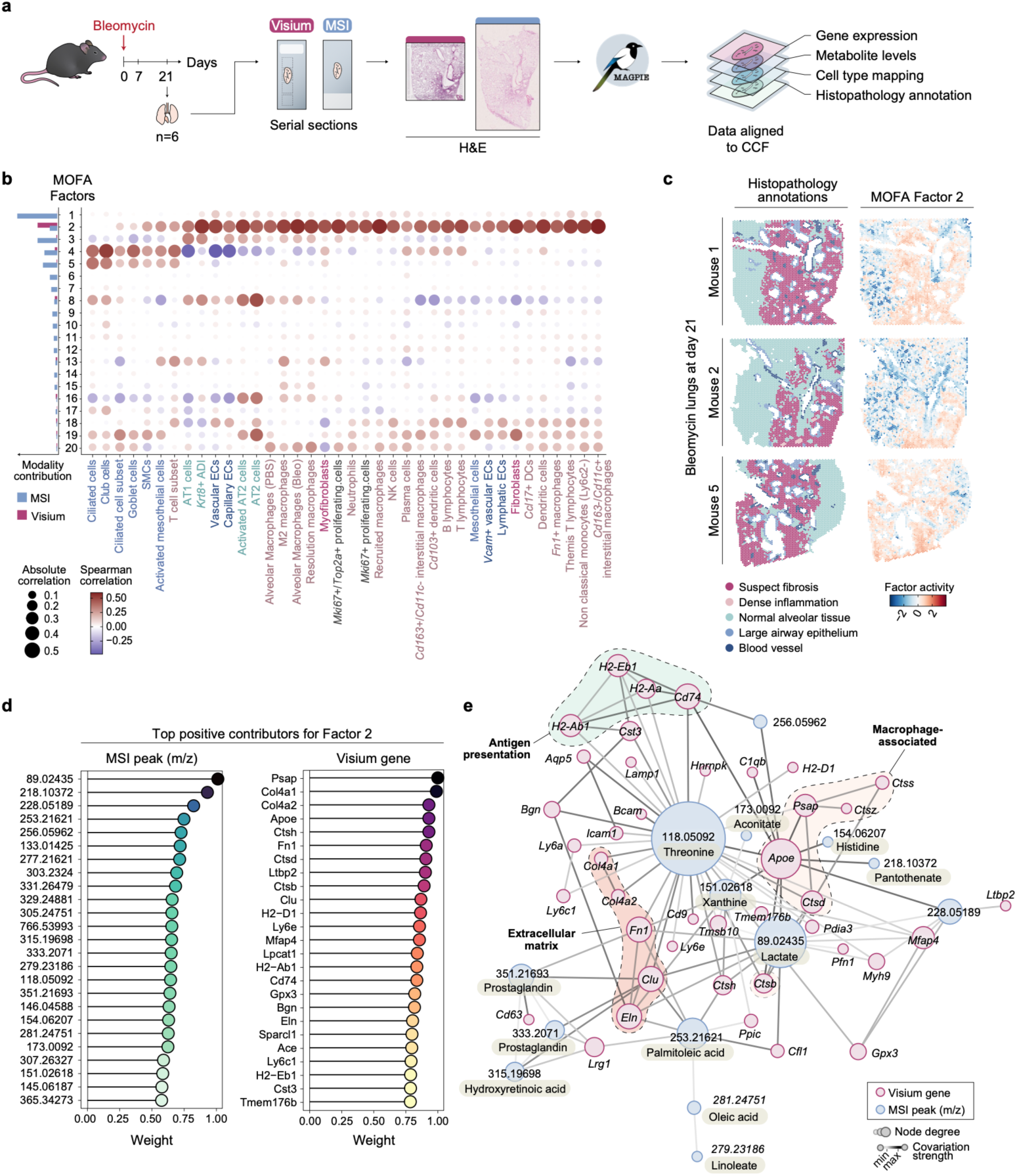
Integrative spatial transcriptomic and metabolomic analysis of bleomycin (BLM)-induced lung fibrosis. **a**, Experimental and computational workflow for generation of spatially resolved transcriptomic (Visium; previously generated^13^ and publicly available data under accession GSE267904) and metabolomic (DESI MSI) datasets from a mouse model of acute BLM-induced pulmonary fibrosis, where lung samples were collected at 21-days following BLM administration. **b**, Multi-omics factor analysis (MOFA)^29,48^ results on aligned Visium and MSI data with 20 factors. Cell type mapping was previously performed using annotated scRNA-seq dataset published by Strunz et al.^47^; inferred cell type densities were correlated with the MOFA factor activities across all spatial locations using Spearman correlation. The relative contribution of modalities for each factor is summarised as a bar chart on the left side. **c**, Spatial plots of the histopathology annotations, based on the Visium H&E images, next to the activity of MOFA Factor 2 for three sections. **d**, Weight of the positive contribution of peaks and genes to Factor 2, showing the top 25 features. **e**, Multi-omics covariation network analysis on top 50 genes and top 25 peaks in Factor 2. The graph was constructed using *GENIE3*^54^ where the top gene-gene, gene-peak and peak-peak edges are selected for the network visualisation. Edge weights (darker edge colour) are proportional to covariation strength from *GENIE3* and node sizes are proportional to node degree. Nodes, representing genes and peaks, are coloured by modality. Abbrev. CCF, common coordinate framework; SMCs, smooth muscle cells; ADI, alveolar differentiation intermediate; ECs, endothelial cells; DCs, dendritic cells.

The co-registration of the modalities into a shared coordinate system enabled us to summarise patterns across modalities using a multi-omics factor analysis (MOFA)^29,48^ for all samples, resulting in 20 factors. These factors were inspected for variance explained, modality contribution, and correlation with cell type density scores^13,47^ (**Fig. 3b**). While Factor 1 had the highest overall variance explained, the contribution was predominantly driven by the MSI modality. Several other factors, e.g. Factors 2, 4, 8, 16, and 19, displayed contributions from both modalities. Factor 2 showed a strong correlation with many disease-relevant cell types, including fibroblasts, myofibroblasts, interstitial/recruited macrophages, and other immune cell populations. Factors 8, 16, and 19 overlapped with alveolar type II (AT2) cells, while Factor 4 was associated with club, goblet, and ciliated cells commonly present in the ciliated epithelium of the lung (**Fig. 3b, Supplementary Fig. 2a**).

Upon spatial inspection, we noted an overlap of areas with high Factor 2 activity and those manually annotated as suspected fibrosis (**Fig. 3c, Supplementary Fig. 2b**). Examination of genes and peaks with the highest contribution to this factor revealed a distinct molecular signature which included a mix of genes associated with extracellular matrix (ECM) components (e.g. *Mfap4, Col4a1, Col4a2, Fn1, Bgn, Eln*), macrophage activity (e.g., *Cd74*, *Apoe*, *Psap*), immune functions (e.g. *H2-Ab1, H2-Eb1, H2-Aa, H2-D1*), and cathepsins (e.g. *Ctsh, Ctsd, Ctsb, Ctss*) (**Fig. 3d**). Several of these genes, such as *Psap*, *Cd74*, *Cd9*, and multiple cathepsins, align with markers previously described for a damage-associated macrophage population implicated in lung injury and fibrosis, where they play a role by modulating tissue responses to lung injury^49^ and suggests immune-modulatory and matrix remodelling roles within the fibrotic regions. These findings were consistent with many of the fibrosis-associated genes detected in our previous Visium data analysis^13^, however, a new dimension of metabolomic information could now be directly associated with the transcriptomic alterations.

The top mass spectrometry peaks contributing to Factor 2 were manually annotated as lactate (*m/z 89.02435*) and pantothenate (*m/z 218.10372*), followed by several long-chain fatty acids (FA) (e.g., *m/z 253.21621*, FA 16:1 palmitoleic acid; *m/z 277.21621,* FA 18:3 linolenic acid; *m/z 305.24751*, FA 20:3 eicosatrienoic acid; and *m/z 303.2324*, FA 20:4 arachidonic acid) and amino acids (e.g. *m/z 118.05092*, threonine; *m/z 146.04588*, glutamate*; m/z 145.06187*, glutamine; and m*/z 154.06207*, histidine) (**Fig. 3d**). Lactate has been repeatedly associated with pulmonary fibrosis, signifying a metabolic shift to anaerobic glycolysis, and may be directly involved in TGF-β-induced myofibroblast differentiation^46,50–52^. Factor 2 was further marked by metabolites of the citric acid cycle (TCA cycle; *m/z 173.0092* aconitate) and glutamine metabolism, which have been implicated in TGF-β-induced myofibroblast activity^53^. The detection of TGF-β-related mechanisms within the fibrotic niche is also supported at a transcriptomic level, where fibrosis-associated TGF-β-signaling was previously reported based on the spatial gene expression data from IPF and BLM-injured lungs^13^.

As the identified genes and metabolomic peaks associated with Factor 2 and regions of fibrosis remained somewhat contained within their separate modalities for this analysis, we aimed to better deduce the connectivity and semantic links between these genes and metabolites. By constructing a covariation network using *GENIE3*^54^, a gene regulatory network comprising both genes and metabolite peaks could be inferred (**Fig. 3e**). Many genes were interconnected based on their similar functions, such as those encoding antigen presentation molecules, ECM-related proteins, as well as genes highly expressed by macrophages. Threonine and lactate metabolites emerged as hub nodes, pointing towards the involvement of their metabolism across a wider range of biological processes. Among them, threonine and lactate displayed covariation with *Apoe*, *Psap*, and *Fn1*, aligning with a previously identified fibrotic macrophage signature^49^. Additionally, lactate has been shown to induce profibrotic genes in macrophages^55^, a finding supported by their spatial covariation in our analysis. Another coregulated group of genes and peaks in the network was annotated to molecules involved in prostaglandin (*m/z 351.21693, m/z 333.2071*) and hydroxyretinoic acid (*m/z 315.19698*) metabolism, which associates with the genes *Lrg1*, *Cd63*, *Clu*, and *Fn1*. The presence of eicosanoids, including various prostaglandins, has previously been reported to be elevated in BLM-induced pulmonary fibrosis^46^ and of relevance for fibrotic progression^56^, making it an intriguing subject for further studies given its co-localisation with an increased gene expression of ECM constituents.

In summary, our *MAGPIE*-enabled integrated spatial multimodal analysis of the BLM-injured mouse lungs both complemented the histopathological assessment of the tissue and further revealed new links between transcriptomic and metabolic alterations occurring during lung fibrosis.

### *MAGPIE* facilitates spatial co-detection of drug substance and drug-induced transcriptional responses

The potential for spatial multimodal analyses extends to many areas of application and, as such, we tested its unique utility in studying transcriptional responses linked to local drug distributions within a tissue. Through Visium and DESI MSI spatial multi-omics data collection on consecutive sections, and subsequent co-registration using *MAGPIE*, we could detect the localisation of compound “AZX” in a rat lung sample following inhaled administration, and explore its impact on the surrounding tissue (**Fig. 4a**). The inhaled administration of AZX, a drug previously in preclinical development by AstraZeneca, has been associated with adverse histopathological lung lesions in animal toxicology studies, thus prompting further exploration of its lung distribution and assessment of unwanted effects *in situ*.

**Figure 4.**
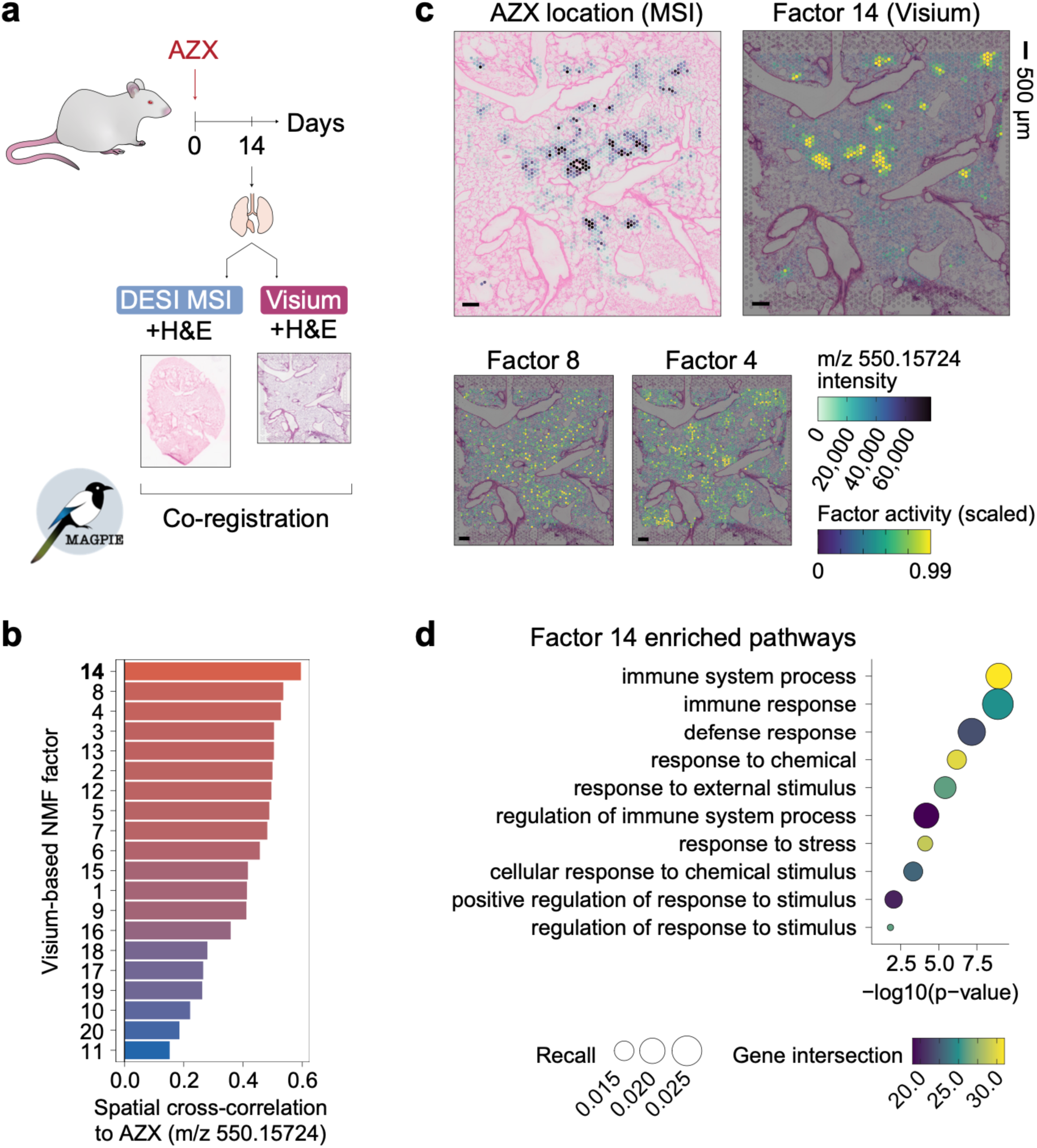
*MAGPIE* co-registration of spatial multi-omics datasets from AZX-treated rat lung. **a**, Outline of experimental design and Visium and DESI MSI data generation from a rat lung exposed to the AZX compound. **b**, Spatial cross-correlation assessment of overlap between MSI AZX compound distribution (*m/z 550.15724*) and Visium NMF factors. **c**, Spatial distribution of MSI AZX intensity distribution (capped at 99^th^ percentile) and the scaled (0-1) Visium NMF factor activities for Factors 14, 8, and 4, overlayed on H&E images. Scale bars corresponding to 500 µm. **d**, Over-representation analysis on the top 100 contributing genes for NMF Factor 14, showing the Gene Ontology (GO) enriched terms. Colours represent gene intersection size between queried genes and genes annotated to each term; dot size corresponds to the recall (ratio between gene intersection size and number of term genes).

Visual inspection of the AZX-dosed tissue revealed that dense AZX hotspots, as detected by MSI, co-localised with variably shaped luminal crystal-like structures observed on microscopic examination of the Visium H&E images (**Supplementary Fig. 3**). After aligning the transcriptome and metabolome data into a shared coordinate space using *MAGPIE*, we investigated the spatial relationship between AZX tissue depositions and local gene expression patterns. The Visium data was first deconvolved into 20 transcriptomics-driven factors using non-negative matrix factorisation (NMF). Next, the spatial cross-correlation between the transcriptomic NMF factors and the intensity of the AZX compound (*m/z 550.15724*), as captured by the MSI modality, could be computed directly owing to the integrated coordinate system. The results identified several transcriptomic NMF factors which showed a spatial association with the compound hotspots, of which Factor 14 showed the highest spatial cross-correlation with the compound, followed by Factors 8 and 4 (**Fig. 4b**). Of these three factors, Factor 14 displayed the most distinct spatially cohesive pattern that corresponded well to the distribution of AZX (**Fig. 4c**) and thus supported a transcriptomic signature localised to areas of drug deposition. Investigation of the underlying Factor 14 transcriptomic signature revealed a profile enriched for immune and defence response pathways, particularly in response to external and chemical stimuli (**Fig. 4d**). This transcriptional signature, identified through the use of integrated Visium and MSI data, suggests a localised inflammatory molecular response triggered by the drug deposition. It further proposes a unique strategy to assess mechanisms of drug-associated toxicity, aiding the development of compounds with a more favourable safety profile.

Here, we demonstrated that multimodal analyses of compound-exposed tissues can provide insights into drug distribution and its associated effects within the tissue. By enabling the integration of joint Visium and MSI datasets within the same spatial coordinate system, *MAGPIE* offers a streamlined approach to correlate pharmacological compound localisation and metabolomic signatures with molecular responses at tissue resolution. This capability of accessible spatial multimodal data generation and computational integration thus holds promise for future studies of disease understanding and drug development.

## Discussion

Through the *MAGPIE* computational workflow, supported by an optimised lung tissue sampling strategy, we have demonstrated the impactful value of integrating spatial transcriptomic and metabolomic datasets to extract insights into disease mechanisms and drug-induced injury in tissues. The *MAGPIE* computational workflow showed versatility across different species and tissue types, as well as MSI technologies, and has been successfully applied to both consecutive section and same-section profiling in human, rat, and mouse datasets. We envision *MAGPIE* to be of high value for users seeking an accessible and generalisable pipeline for integrating spatial multimodal data.

*MAGPIE* allows users the flexibility to incorporate external information and optimise hyperparameters to suit their available samples. Based on our input parameter tests and outcomes, we offer several recommendations for future datasets processed with *MAGPIE* for spatial multimodal integration, particularly when dealing with serial sections. Obtaining a microscopy image from the MSI samples, preferably using the same H&E staining technique as for Visium, is important for accurate landmark identification between MSI and Visium images. High structural similarity between the matching tissue sections should be prioritised whenever possible, especially when consecutive or more distant sections are used for the two modalities. Our examples highlight the need for careful consideration of feasibility before attempting co-registration and that accurate co-registration may require a greater number (≥ 10) of landmarks and/or a non-linear transformation, depending on the similarity between sections. We advise careful assessment of section similarity and a comparison of different co-registration options when producing new datasets.

Moreover, we illustrate the unique benefit of obtaining and integrating paired spatial multi-omics data for studies of disease characteristics as well as local tissue responses to drug depositions. In the BLM mouse model, the integration of spatial transcriptomics and metabolomics revealed distinct signatures associated with fibrotic regions. These alterations were characterised by an increased transcriptionally-inferred presence of inflammatory cells, fibroblasts, and myofibroblasts, coupled with a shift in local metabolism towards states previously described to be relevant for pulmonary fibrosis^46,51,52^. Additionally, the integration of Visium data with MSI-derived information on compound deposition in the lung using *MAGPIE* enabled the identification of an inflammatory transcriptomic signature localised to areas of drug accumulation in the rat lung, potentially providing mechanistic insight into the observed lung toxicity of this compound.

Histology plays a valuable role when interpreted in tandem with SRT and MSI by providing essential structural context for analyses. As the field of spatial omics moves towards establishing a stronger link to histological assessment, preserving tissue morphology and avoiding freezing artefacts becomes critical for accurate pathological interpretation. In this context, formalin-fixed paraffin-embedded (FFPE) tissue samples offer an attractive alternative to fresh frozen tissue. While workflows for Visium and other SRT methodologies have been established for FFPE tissues, integrating MSI with FFPE samples requires careful consideration of sample preparation protocols to ensure preservation of chemical identity and spatial localisation, particularly for smaller metabolites^57–60^. As these protocols continue to develop, we expect that spatial multimodal datasets using both fresh frozen and fixed tissues will become increasingly important for spatial studies investigating molecular mechanisms in tissue biology. Due to the flexibility of the pipeline, we anticipate that the *MAGPIE* workflow will be adaptable to datasets produced with different tissue preparation methods and Visium platforms. Although the *MAGPIE* pipeline is designed to utilise the standard file format for spatial data introduced by Visium, the input format of the second modality is a simple tabular matrix containing features and spatial coordinates. Therefore, it may be further applicable to measurements from other omics beyond MSI as more combinations of modalities become possible. For example, as co-registration with images from the same tissue section is already supported by the *MAGPIE* pipeline, it is readily applicable to combined assessment of metabolomics, transcriptomics, and proteomic data, e.g. from multiplex immunofluorescence imaging. The co-registration of spatial transcriptomics with spatially resolved proteomics or spatially resolved chromatin accessibility profiling^61,62^ would enable a greater understanding of the dynamics of gene regulatory relationships in space. This could be achieved through inference of gene regulatory networks using computationally combined observations across modalities, by adapting single-cell methods^63–65^ or using novel spatially resolved methods as the analysis landscape continues to develop.

In conclusion, we have shown that *MAGPIE* is a robust, versatile, and reproducible pipeline for the co-registration and computational integration of spatial transcriptomic and metabolomic data, which enables richer biological insights from tissues, showcased by its application to characterise molecular responses to tissue injury. By continuing to refine and expand spatially resolved analysis techniques, and in parallel developing new computational approaches for processing and analysing the data, we can further enhance the resolution and applicability of spatial multi-omics analyses in biomedical research and enable the generation of previously unattainable molecular insights into human disease for the discovery of novel treatments.

## Supplementary Figures

**Supplementary Figure 1.**
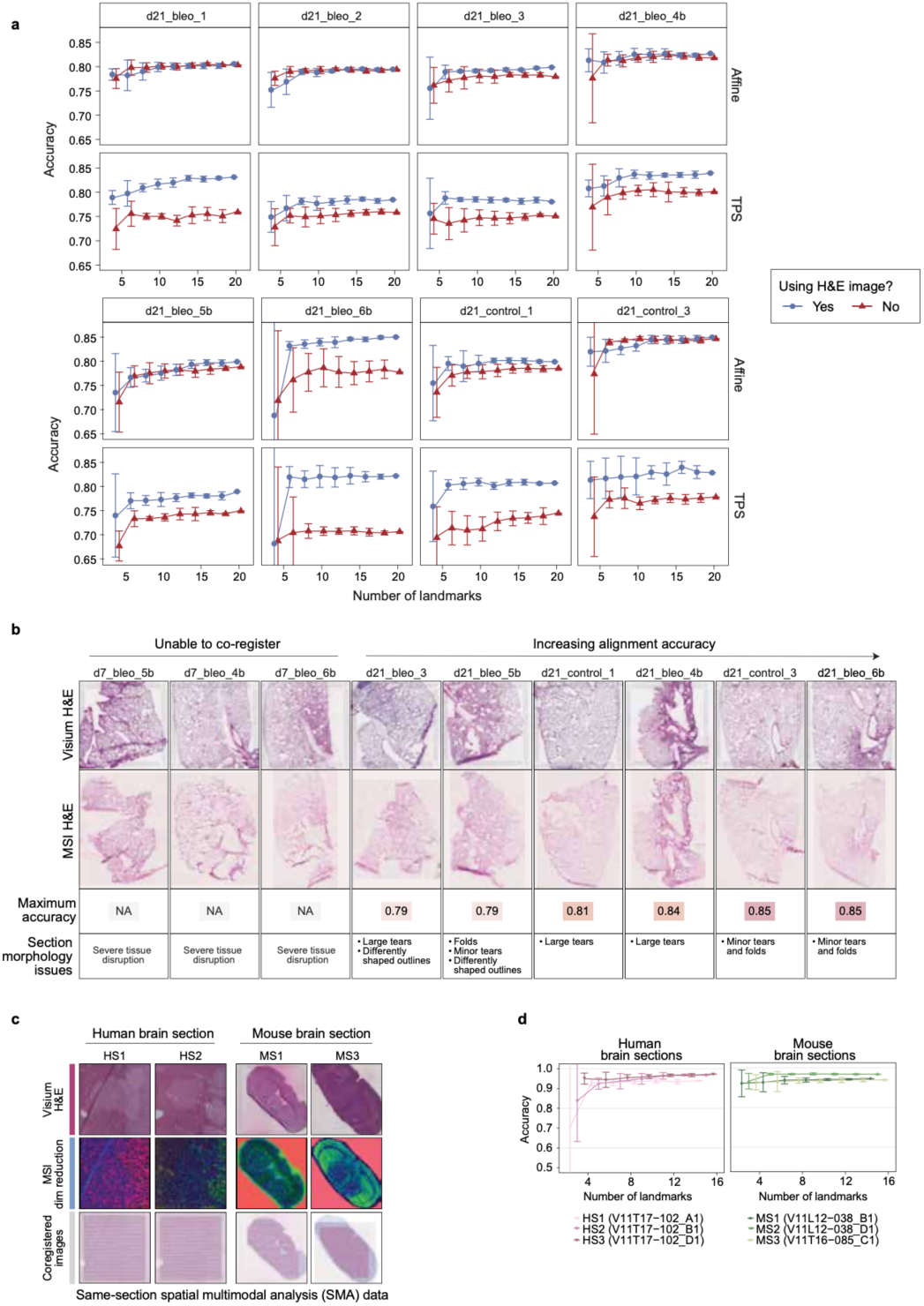
Further assessment and benchmarking of the flexibility and robustness of the MAGPIE integration pipeline. **a**, Evaluation of varying *MAGPIE* hyperparameters in the BLM mouse model dataset showing alignment accuracy scores of tissue-to-background matching between Visium and MSI data after co-registration across a range of landmarks, employing a linear (affine) and non-linear (TPS) transform, and with or without using an intermediate MSI H&E image to assist with co-registration. Mean and 95% confidence intervals of accuracy (y-axis) are shown across 5 repeats for 4-20 landmarks (x-axis) and the use of an intermediate MSI H&E image is shown through the colour and shape. **b,** Ranking of the BLM mouse model samples by similarity (maximum accuracy measurement) between Visium and MSI sections comments on tissue section morphology artefacts that may influence alignment accuracy. **c,** Illustration of flexibility of MAGPIE across different species and tissue types by showcasing its application to all samples in same-section Visium and MSI data from mouse and human brain tissue, followed by co-registered coordinates from both modalities. **d,** Variation in accuracy of tissue against background matching between Visium and MSI data after co-registration (no intermediate MSI image provided) across a range of landmarks (3-15) with 5 repeats each. Error bars correspond to 95% confidence intervals.

**Supplementary Figure 2.**
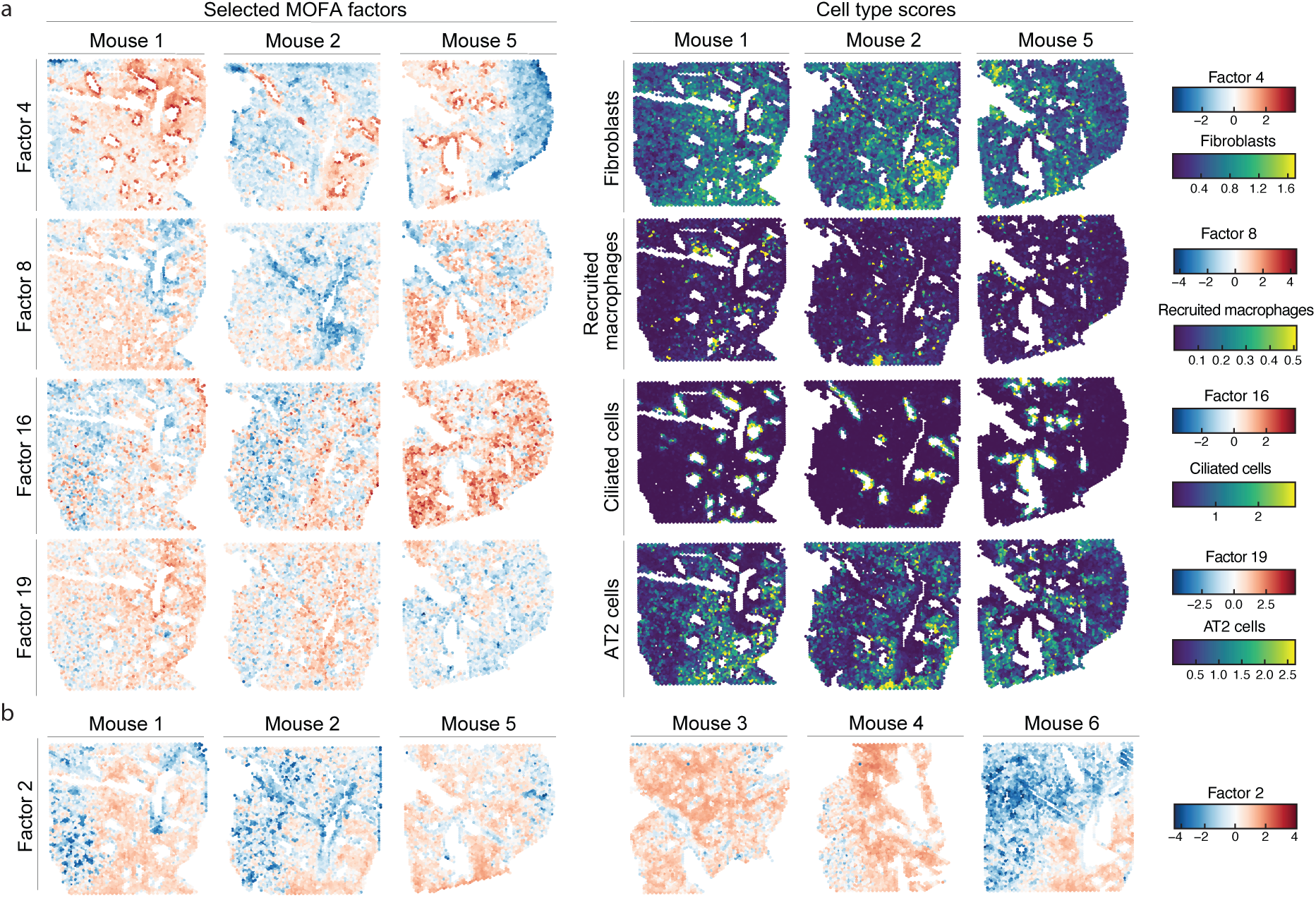
Spatial distribution of MOFA factors and key cell types in a bleomycin-treated mouse model. **a**, Spatial distributions of selected MOFA factors that showed contribution from both modalities (left) and key cell types which showed correlation with the shown MOFA factors and with MOFA Factor 2 (right, capped at 99th percentile). **b,** Spatial plots of the activity of MOFA Factor 2 across all 6 included samples.

**Supplementary Figure 3.**
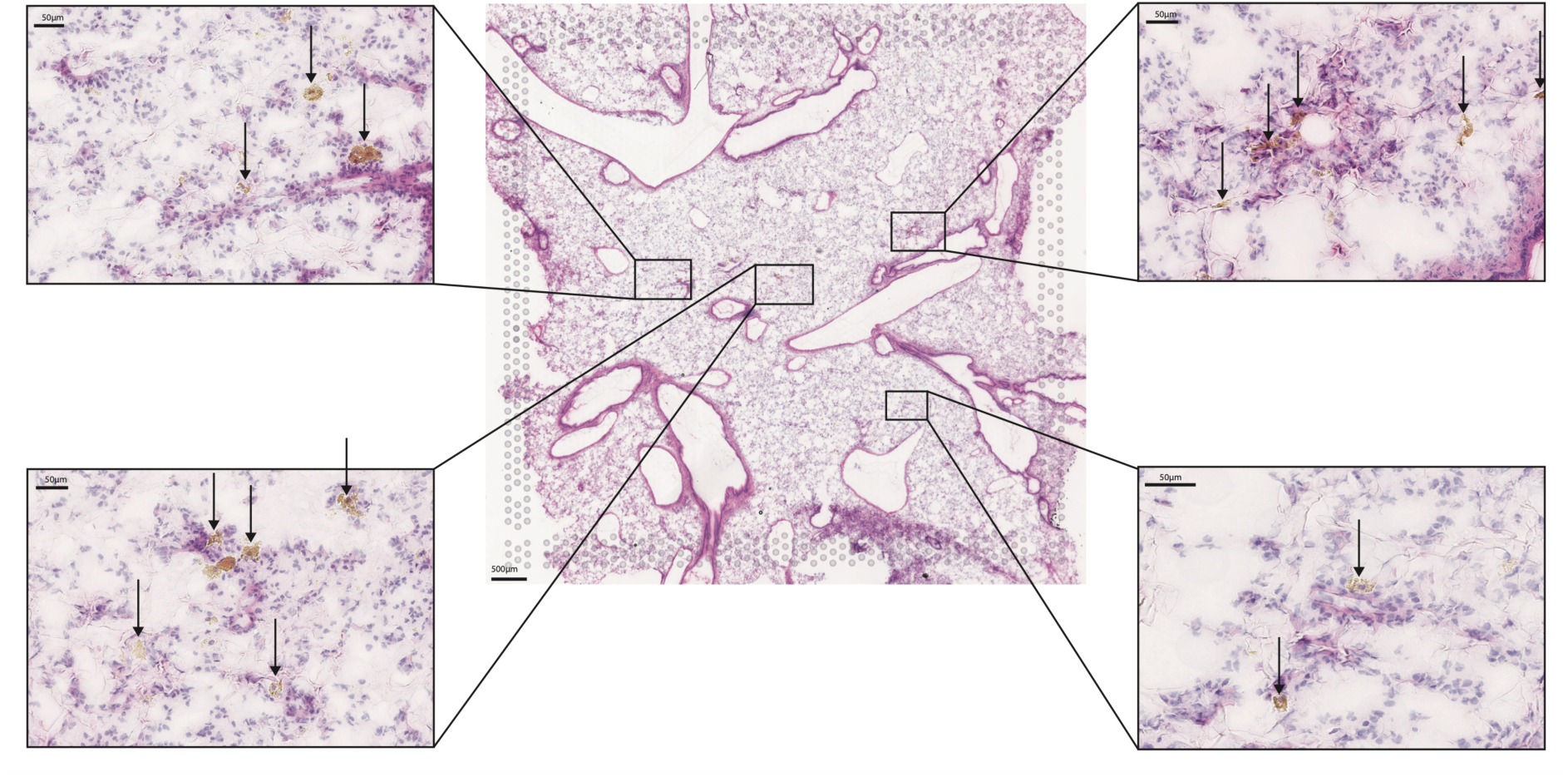
Detection of brown-coloured crystal-like structures (black arrows), co-localising with dense AZX deposits, in a compound-treated rat lung sample, as seen in the H&E-stained Visium sections. Centre image scale bar indicates 500 µm. Scale bars in the zoomed-in inserts measure 50 µm.

## Methods

### Experimental data generation

#### Ethical considerations

Animal care and handling adhered to the standards established by the Council of Europe ETS123 AppA, the Helsinki Convention for the Use and Care of Animals, Swedish legislation, and AstraZeneca global internal standards. All experiments were ethically approved by the Gothenburg Ethics Committee for Experimental Animals in Sweden, complying with Directive 2010/63/EU. The studies received local Ethical committee approval in Gothenburg (EA000680-2017 and 2020-002853) with the assigned site number 31-5373/11.

#### Animals

##### Bleomycin-induced lung fibrosis mouse model

The mouse lung sample generation and collection were previously described in the study by Franzén and Olsson Lindvall et al.^13^. In short, female C57BL/6NCrl mice (Charles River, Germany; eight weeks of age upon arrival) were housed in a facility with ad libitum access to food (R70, Lantmännen AB, Vadstena, Sweden) and tap water. Following a five-day acclimatization period, mice were subjected to oropharyngeal administration of either 30 μl of bleomycin (Apollo Scientific, BI3543, Chemtronica Sweden; 40 µg/mouse) dissolved in saline or saline alone (vehicle control). Lung samples were collected on day 7 (d7) and day 21 (d21) post-bleomycin challenge, capturing both the early inflammatory and tissue remodelling phase (d7) and established tissue damage (d21).

##### AZX-dosed rat lung

Male Wistar Han rats (Charles River, Germany) were seven weeks old upon arrival and approximately 10 weeks at the start of dosing. They were group caged, 2-6 rats per cage, and maintained on a normal day/night cycle of dark 18:00 and light 06:30 at a uniform temperature (>25°C) and humidity (<5%), with access to chew sticks and nesting material. Rats had free access to diet (R70) and tap water. Animals were randomised and regrouped into new cages prior to the first dose and housed with others in the same dosing group to minimise contamination by grooming and coprophagia. The animal used in this study (n=1) was subjected to daily inhalation exposure via snout-only administration for 14 days with 15 mg/kg/day of the AZX test formulation consisting of AZX (10%), MPEG-2000-DSPE (1.25%), trileucine (5%), and trehalose (83.75%), in a flowpast exposure chamber. The animal was acclimated to the method of restraint over a five-day period preceding the first test substance exposure. Animal body weight was recorded prior to the study, twice weekly, pre-dose, and at termination.

#### Tissue processing for spatial multimodal analysis of rodent lung

The rodent lung tissue sampling protocols were optimised to enable high quality analysis using both the Visium Spatial Gene Expression platform and mass-spectrometry imaging (MSI), both with regards to analyte preservation and histological integrity. Fresh frozen tissue specimens are typically embedded in optimal cutting temperature (OCT) compound for Visium analysis, however, OCT is known to interfere with the quality of mass spectrometry readouts and should therefore be avoided or removed in MSI experiments^21,22^. Thus, we opted for careful inflation of the lungs using a low-melting point agarose solution prior to snap freezing of the tissue without any further embedding material.

For the mouse study, the mice were first anaesthetised using isoflurane (5% concentration, air flow ∼2 L/min) and maintained with 3% isoflurane (air flow ∼0.7 L/min). A midline incision was made from the abdominal midsection to the chin. Subsequently, 0.1 mL of heparin was injected through the diaphragm into the heart, followed by severing the abdominal aorta to exsanguinate the mice. The heart and right lung lobes were tied off and removed. The pulmonary circulation was perfused with 37°C saline followed by 37°C low-temperature melt agarose (SeaPlaque) solution. The lung was then gently inflated with 0.4-0.5 mL of 37°C agarose solution via the trachea and tied off. The lung tissue was collected and snap-frozen in pre-chilled NaCl over dry ice and stored at −80°C for further analyses.

For the rat sample, the trachea was surgically exposed and secured using a 2-0 silk suture on the day of termination. Subsequently, 0.4 mL of heparin was injected into the heart through a small incision in the diaphragm, after exposing the abdominal wall. The pulmonary circulation was flushed via the pulmonary artery using a solution of 15 mL of 37°C saline followed by 7 mL of 37°C low-melting point agarose (SeaPlaque) solution (0.75 g agarose in 50 ml saline). A suture was placed around the heart to prevent heart leakage. A cannula was then introduced into the trachea and the lung was gently inflated with 4.5 mL of agarose solution. Lungs and heart were removed as a single unit, placed in ice-cold PBS, and allowed to rest on wet ice for approximately 20 minutes to solidify the agarose. The right superior lobe of the lung tissue was thereafter placed on pre-chilled tin foil in a plastic fixation cassette. The sample was rapidly frozen by immersion in isopentane, pre-chilled in liquid nitrogen. Samples were stored at −80°C until further analysis.

#### Spatial transcriptomics and metabolomics experimental procedures

##### Sectioning and RNA assessment

Agarose-inflated mouse and rat lung tissues were mounted to the cryostat specimen chuck using a drop of purified water, and thereafter cryo-sectioned at 10 µm thickness with the cryostat chamber temperature set to −20°C and −10°C for the specimen head. It is important to note that the frozen tissue samples are never embedded in OCT compound, given its incompatibility with MSI analysis. Consecutive sections from each lung were thaw-mounted on a Visium slide for spatial transcriptomics and a Superfrost slide (Fisher Scientific, Loughborough, UK) for desorption electrospray ionization (DESI) MSI. Slides were stored in −80°C until further analyses. For RNA quality assessment, approximately ten sections from each lung tissue were collected and stored at −80°C prior to RNA extraction using the RNeasy micro kit (Qiagen). RNA quality was assessed using a 5300 Fragment Analyzer (Agilent) and the RIN values were >8 for all samples.

##### Generation of spatially resolved transcriptomics data

For the AZX-treated rat sample, tissue fixation and staining followed the Methanol Fixation, H&E Staining, and Imaging Visium protocol (10X Genomics). Imaging was performed at 40X magnification using the Aperio Digital Pathology Slide Scanner (Leica Biosystems) and sequencing libraries were prepared according to the Visium Spatial Gene Expression User Guide (10X Genomics; Rev E). The rat library (n=1) was sequenced on the NovaSeq 6000 (Illumina) platform with an S4 flowcell using the following set-up: Read1: 28 bp, Index 1: 10 bp, Index 2: 10 bp, Read2: 90 bp. A 1% PhiX spike-in was included in each run. A total of 177 M reads was generated for the rat sample.

##### Desorption electrospray ionisation mass spectrometry imaging (DESI MSI)

For both mouse (n=11) and rat (n=1) lung samples, DESI MSI was carried out using an automated 2D DESI source (Prosolia Inc, Indianapolis, IN, USA) with a home-built sprayer assembly mounted to a Q-Exactive FTMS instrument (Thermo Scientific, Bremen, Germany). Analyzes were performed at spatial resolutions of 65 μm in negative ion mode and mass spectra were collected in the mass range of 80−900 Da with mass resolving power set to 70000 at *m/z* 200 and an S-Lens setting of 100. Methanol/water (95:5 v/v) was used as the electrospray solvent at a flow rate of 1.0 μL/min and a spray voltage of −4.5kV. Distance between DESI sprayer to MS inlet was 7mm, while distance between sprayer tip to sample surface was 1.5mm at an angle of 75°. Nitrogen N4.8 was used as nebulizing gas at a pressure of 6.5 bar. Omnispray 2D (Prosolia, Indianapolis, USA) and Xcalibur (Thermo Fisher Scientific Inc) software were used for MS data acquisition. Individual line scans were converted into centroided .mzML format using MSConvert (ProteoWizard toolbox version 3.0.4043) and subsequently into .imzML using imzML converter v1.3. Haematoxylin and eosin (H&E) staining was performed post-analysis on the same tissue sections and the stained sections were imaged at 20x with Aperio CS2 digital pathology scanner (Aperio Tech., Oxford, UK), and visualised with QuPath 0.23^66^ for histopathological annotations performed by a pathologist (L.S.).

### Computational data processing

#### Preprocessing of mass spectrometry data

For both the bleomycin mouse model and AZX-treated rat datasets, the .imzml data files were initially imported into SCiLS Lab software (Bruker Daltonics, Germany, 2022a MVS). Bisecting k-means segmentation was performed within SCiLS Lab to create two regions of interest (ROIs): tissue and background (area outside the tissue). Individual samples were then segmented using the tissue/background ROIs with some manual alterations. Individual pixel-level total ion count (TIC) normalised peak intensities were extracted using the SCiLS Lab API, which yields a pixel-by-peak table with associated metadata. This table was later used as input to the *MAGPIE* pipeline. The peaks were annotated against compounds in the KEGG database^67^ using [M-H]^-^ adducts with a mass tolerance of 5 ppm. Only peaks which could be annotated to a metabolite were retained for the downstream analysis of the bleomycin-treated mouse samples.

#### Preprocessing of Visium data

Raw FASTQ files were processed with Space Ranger 1.2.2 (10x Genomics). Sequencing reads were aligned to their respective reference genomes: rn6 for the rat sample and mm10 for the mouse samples. Alignment of H&E images to the fiducial frame was performed manually using the Loupe Browser software (v.6, 10X Genomics). For the mouse study, the Space Ranger output data and associated metadata, including cell type deconvolution results, was published in conjunction with the publication by Franzén and Olsson Lindvall et al.^13^, and can be downloaded from Gene Expression Omnibus (GEO) (accession number GSE267904) and BioStudies (S-BSST1409).

### The *MAGPIE* pipeline

The *MAGPIE* pipeline is written in Python (v3.10.11) and presented as a Snakemake workflow, with dependencies on *snakemake*, *shiny*, *matplotlib*, *pandas*, *numpy*, *scikit-image*, *pathlib*, *scikit-learn*, *scipy*, *json*, *collections*, *shutil*, *gzip*, *h5py*, and *scanpy*. All pipeline components can be initiated from the command line; detailed documentation for each component is provided at https://core-bioinformatics.github.io/magpie. The full pipeline is open source under the MIT/ CC-BY licence and is found on GitHub at https://github.com/Core-Bioinformatics/magpie.

As input, the *MAGPIE* pipeline takes the Visium data in the standard Space Ranger output format (including *filtered_feature_bc_matrix.h5* containing gene expression information, *tissue_hires_image.png* with the H&E image, *tissue_positions.csv* containing spatial coordinates per spot and *scalefactors_json.json* translating from H&E image to spatial coordinates) and a peak-by-pixel table for the MSI data, with the option to add a microscopy image for the MSI section. The image alignment within *MAGPIE* utilises a landmark-based approach. Users may opt to select their own landmarks outside the pipeline, either manually or automatically using an external tool (e.g. ELD^23^). *MAGPIE* includes an interactive Python Shiny application which can be used before the Snakemake pipeline. Within this application, a lower dimensionality image is created based on the MSI data using a user-selected method, for example, using the first or first 3 principal components (based on all or a subset of peaks) as colour channels or using an individual peak of interest based on *a priori* knowledge. For manual landmark identification, users are thereafter prompted to select landmarks within the interactive application, either (1) between MSI dimensionality reduction image and Visium H&E image or (2) between MSI dimensionality reduction image and MSI microscopy image, if available, then between the MSI microscopy image and Visium H&E image. If users identify landmarks externally, they may add them in a tabular format and skip the in-built landmark selection tool.

The next stage of the pipeline is streamlined within a Snakemake workflow, which consists of three main modules. In the first module, the identified landmarks are used to map MSI coordinates to Visium coordinates, using either a linear (affine) or non-linear (TPS) transform based on the scikitimage implementation (v0.24) in accordance with user selection. This results in a common coordinate framework (CCF) between the two modalities where each (x,y) coordinate in the MSI modality can be directly mapped into the Visium modality. The second module of *MAGPIE* prepares and stores the new MSI data in a Space Ranger-style object, including a .h5 file and a ‘spatial’ folder containing coordinates and images, which can be read by other spatial analysis ecosystems. As a third optional module, the data points for the MSI modality can be aggregated into the Visium spot barcodes, creating a 1:1 spatial map between the MSI and Visium observations, following a strategy described in the next section.

#### Aggregation of MSI data per Visium spot

For performing fully integrated spatial multimodal analysis, we match MSI pixels to Visium spots to create combined ‘observations’. Initially, we calculate the between-spot distance in the Visium data then find the MSI pixels within half that distance for each Visium spot (the spots’ “expanded radius”) (**Fig. 1c**). This is calculated efficiently by finding the k-nearest neighbours for each spot and then selecting the neighbours within the specified expanded radius distance. MSI pixels within each expanded Visium spot are then either averaged or summed (user-defined option) to create the new matching observations. This step of the data integration was implemented within the *MAGPIE* Snakemake workflow as a final optional module and as a separate function, “CreateMultiModalObject”, in the *semla* R package (v. ≥ 1.3.0)^26^, with which the loaded Visium and *MAGPIE*-aligned MSI data can be joined into a multimodal object.

### *MAGPIE* pipeline benchmarking

#### Hyperparameter testing

The performance of co-registration was assessed on the overlap between MSI and Visium observations after coordinate transformation. We relied on the tissue/background annotation identified through the Space Ranger pipeline and applied tissue/background labels to the MSI data based on the first principal component, using data-specific thresholds identified using underlying microscopy and lower dimensional images. We then calculated summary statistics to capture the accuracy of the transformation to match tissue to tissue and background to background (i.e. # spots annotated as tissue in both modalities + # spots annotated as background in both modalities divided by total # spots). For each sample (n=8) in the bleomycin-treated mouse dataset, 20 landmarks were identified (1) between MSI dimensionality reduction and MSI H&E then between MSI H&E and Visium H&E and (2) between MSI dimensionality reduction and Visium H&E directly. For numbers of landmarks ranging between 4 and 20 (with 5 repeats), the given number of landmarks was sampled from the total pool, the transformed coordinates calculated and pooled into matching Visium/MSI spots and the alignment accuracy of tissue/background labelling recorded. Samples were ranked for image similarity based on the best accuracy achieved across all numbers of landmarks and repeats.

#### Same-section spatial multimodal data

For the same-section Visium MALDI MSI multimodal data tests, the datasets were acquired from Mendeley Data (DOI: 10.17632/w7nw4km7xd.1) and three mouse brain (V11L12-038_B1, V11L12-038_D1, V11T16-085_C1) and three human brain (V11T17-102_A1, V11T17-102_B1, V11T17-102_D1) datasets were selected. A total of 15 landmarks were identified per sample and a number of landmarks ranging between 3 and 15 (with 5 repeats) was tested using the same approach as described above.

#### Downstream tissue injury study multimodal data analysis

##### Bleomycin mouse model of pulmonary fibrosis

MAGPIE-aligned MSI data, using TPS transformation^68^, for mouse lung samples collected 21 days after bleomycin administration (one tissue section per animal; n=6) was loaded into R with *semla*. The corresponding Visium Space Ranger output data files (GSE267904^13^) were furthermore imported with “ReadVisiumData” where spots within the tissue and image borders were kept. The Visium data was filtered to exclude genes starting with “Rpl”, “Rps”, “Mrp” and “mt-“, pseudogenes and noncoding genes and focus solely on protein-coding genes detected in > 5 spots. Spots were further filtered to remove those with 100 or fewer transcripts and 10% or higher mitochondrial or haemoglobin gene content. The MSI data was TIC-normalised beforehand, and once in a *semla* object format, it was filtered to keep pixels with more than 300 detected peaks and a total intensity of > 14×10^5^. Peaks that had not been annotated to metabolites (using M-H adduct) were excluded from the downstream analysis. Further, peaks with the same mass down to two decimal points were combined by taking the peak with the highest sum of intensities across all samples. A multimodal *semla* object was thereafter created with “mean” as the aggregating function to obtain average m/z peak intensity values per Visium spot. The full Visium data was subsequently normalised with “NormalizeData” and scaled with “ScaleData”. All annotated MSI peaks (scaled per peak) and top 2000 most abundant genes were used to create a MOFA+^29^ model with 20 factors (medium convergence mode). Cell type deconvolution results were obtained from the previously generated data by Franzén and Olsson Lindvall et al.^13^; the Spearman correlation was calculated between inferred cell type densities and MOFA factor activities across spots. Where factors showed the highest correlation with any cell type in the negative direction, the factor activity scores were reversed to favour interpretability (specifically factors 1, 2, 3, 4, 8, 9, 10, 12, 14, 15, 18, 19, 20). Spot factor scores were compared to histopathology annotations and the top genes and peaks in factors showing clear overlap with fibrotic and inflamed regions were inspected. Multi-omics covariation network analysis was performed on the top 50 genes and 25 peaks in Factor 2 using *GENIE3*^54^. The Spearman correlation between all peaks and genes was also calculated and connections between nodes which showed a negative correlation were removed to ensure all edges corresponding to positive covariation. The top gene-gene, gene-peak, and peak-peak edges were selected (proportional to number of nodes included for each modality, specifically top 40 for gene-gene and gene-metabolite edges and top 20 for metabolite-metabolite) and the resulting network was visualised using the R package *igraph*, with the Davidson-Harel layout algorithm using cooling factor 0.8. Disconnected subgraphs containing ≤ 3 nodes were removed from the network. Edge weights are proportional to covariation strength from *GENIE3* and node sizes are proportional to degree in the network.

##### AZX-dosed rat lung tissue

After aligning observations between datasets, as detailed above, data from each modality was separately loaded into a *semla*^26^ object, removing spots outside of the tissue and image area. The Visium data was filtered to remove genes matching the regex patterns “^Rpl|^Rps”, for ribosomal genes and “^Mt-“, for mitochondrial genes, genes annotated as non-coding or pseudogenes, as well as genes detected in 5 or fewer spots. Spots with ≤ 300 transcripts or ≥ 30% mitochondrial or haemoglobin-related gene content were further removed to exclude low-quality spots. The TPS-aligned and TIC-normalised MSI data was filtered to keep only peaks with a metabolite annotation and *m/z 550.15724* (AZX compound), and pixels with more than 300 assigned peaks and a total (normalised) ion intensity measure exceeding 23×10^5^ were kept. The joint multimodal *semla* object was thereafter created using the function “CreateMultiModalObject”, with the data aggregation argument set to “mean” to group the MSI data into the Visium coordinates by averaging the peak intensities. Normalisation and scaling of the Visium data were thereafter performed using the Seurat functions “NormalizeData” and “ScaleData”. NMF analysis, specifying 20 factors, was run on the Visium data using the “RunNMF” function from the *singlet* R package^69^. Spatial co-localisation with m/z 550.15724 in the metabolomics modality was assessed using the spatial cross-correlation metric from *MERINGUE*^70^. Over-representation analysis of Gene Ontology (GO) biological processes terms was performed on the 100 top contributing genes for NMF Factor 14 using the R package *gprofiler2*^71^, with the organism set to “rnorvegicus”.

## Code availability

The *MAGPIE* source code is publicly available at https://github.com/Core-Bioinformatics/magpie. All the code used to perform the downstream analyses will be available upon publication at GitHub and Zenodo.

## Data availability

The previously generated Visium data from the bleomycin mouse model is available in Gene Expression Omnibus under accession number GSE267904; all associated metadata is available at BioStudies with accession number S-BSST1409. The mouse and human brain tissue samples from the study by Vicari et al. were downloaded from Mendeley Data, DOI: 10.1038/s41587-023-01937-y. The new data generated in this study will be available upon publication.

## Acknowledgements

This work was supported by the Swedish Foundation for Strategic Research (grant no. ID18-0094 supporting L.F.), the MRC DTP iCASE studentship programme (G117817 supporting E.C.W.), and AstraZeneca. IM was funded by the Wellcome Trust [203151/Z/16/Z] and the UKRI Medical Research Council [MC_PC_17230]. We thank A. Collin and E. Sand for assistance and input on tissue sample collection and assessment, A. Borde and T. Volckaert for providing the mice from the bleomycin mouse model study, B. Keith for Visium data pre-processing, M. Hühn and V. Ptasinski for initial results discussions, and R. Mauron, M. Machado, and M. Ekvall for input on multimodal integration. For the purpose of open access, the authors applied for a CC BY public copyright license to all versions of the manuscript arising from this submission.

## Author information

### Contributions

E.C.W., L.F., JT and IM designed and implemented the pipeline and performed downstream analysis. J.D. and A.H. provided support for early versions of the co-registration pipeline. J.E.M. contributed to the design of the data mapping strategy. S.O. optimised the rodent lung sampling strategy. J.J.H., A.O., M.S., L.F, M.O.L., and M.M.M planned the experimental work. M.O.L. sectioned the rodent lung tissues and generated the rodent Visium data. M.M.M. provided the rat lung samples. G.H. ran the MSI experiments for the rodent lung samples and G.H., E.C.W. and A.Z. performed the MSI data pre-processing. L.S. performed the histopathological assessment of the rodent lung tissues. E.C.W. and M.V. performed manual landmark selection of samples. J.E.M., M.V., and J.L. contributed to methodological discussions. Data interpretation was done by L.F., M.O.L., E.C.W. and G.H. L.F. and E.C.W. created the final figures and illustrations. E.C.W., L.F., M.O.L., I.M. wrote the manuscript with input from other authors. J.J.H., A.O., P.L.S., M.S., J.T. and I.M. guided and supervised the project.

